# Credit Assignment via Behavioral Timescale Synaptic Plasticity: Theoretical Frameworks

**DOI:** 10.1101/2025.06.12.659336

**Authors:** Ian Cone, Claudia Clopath, Rui Ponte Costa

## Abstract

Behavioral Timescale Synaptic Plasticity (BTSP) is a form of synaptic plasticity in which dendritic Ca^2+^ plateau potentials in hippocampal pyramidal neurons drive rapid place field formation. Unlike traditional learning rules, BTSP learns correlations on the timescales of seconds and rapidly changes single-unit activity in only a few trials. To explore how BTSP-like learning can be integrated into network models, we propose a generalized BTSP rule (gBTSP), which we apply to unsupervised and supervised learning tasks, in both feedforward and recurrent networks. Unsupervised gBTSP mirrors classical frameworks of competitive learning, learning place field maps (in the feed-forward case), and attractive memory networks (in the recurrent case). For supervised learning, we show that plateau events can reduce task error, enabling gBTSP to solve tasks such as trajectory matching and delayed non-match-to-sample. However, we find that credit assignment via gBTSP becomes harder to achieve with increased network depth or CA3-like recurrence. This suggests that additional features may be needed to support BTSP-mediated few-shot learning of complex tasks in the hippocampus.

## Introduction

Recent experimental observations have revealed the existence of a novel plasticity phenomenon occurring in hippocampus, termed “Behavioral Timescale Synaptic Plasticity” (BTSP). BTSP occurs in hippocampal pyramidal cells following strong, dendritic “plateau potentials”, and has been observed to be a “primary” driver of field formation (such as place fields) in both hippocampal areas CA1^1–3^ and CA3^4^. Unlike other established learning rules, BTSP operates over a wide temporal range, potentiating and depressing inputs which were active seconds before or after a postsynaptic “plateau” event. A similarly distinctive feature of BTSP is its rapid learning speed - once triggered, it can form long-lasting hippocampal fields in a one-shot or few-shot manner.

While the discovery of BTSP has advanced our understanding of single-cell learning in hippocampus, we are still lacking a comprehensive theoretical framework to understand how BTSP may contribute to network level plasticity. For example, experiments have shown that inputs from entorhinal cortex layer 3 (EC3) are necessary to trigger plateau potentials in CA1, and as such, have been hypothesized to act as a sort of “target signal”^5^ which guides plateau generation. But what sort of “targets” should EC3 produce? That is, “when” and “where” should BTSP-triggering plateau events occur for hippocampal learning to be successful? Or, from a more general perspective, “when” and “where” should BTSP events occur to optimize a network’s function?

Furthermore, how can we reconcile the hallmarks of BTSP (wide temporal kernel and few-shot learning) with traditional learning frameworks (particularly supervised ones)^6–13^, for which small learning rates and temporally precise credit assignment are generally required for convergence? Does the presence of BTSP put constraints on the possible tasks and the neural architectures (i.e. circuit designs) which can learn them?

Previous theoretical work on BTSP has described in detail its single-cell properties^1,2,14^ and consequences in memory networks^4,15,16^. However, such work has thus far have been forced to assume specific, hand-tuned plateau induction protocols. In contrast, this work aims to formulate BTSP in such a way that we can describe where and when post-synaptic plateaus should occur such that the network learns a given unsupervised or supervised objective.

Towards this goal, we formulate a generalized BTSP rule which can a) match existing experimental data, and b) give us analytical, differentiable expressions for how the learning performance (i.e. the loss) depends plateau events. We take this rule and demonstrate its ability to learn in both feed-forward and recurrent networks, on both supervised and unsupervised tasks. Since we derive an analytical expression for plateau “function”, we can predict the occurrence of plateau events, given that we know the weights, inputs and task. Further, by applying constraints on our expression for plateau events, we can approximate the sparse nature of these events in vivo. However, we show that the rapid, one-shot formation of single fields associated with BTSP runs into critical stability issues when applied in deep/recurrent networks, because of exploding and vanishing mathematical terms. We finish by discussing how potential architectures and tasks may be able to avoid this issue, and what the implications are for our understanding of hippocampal networks.

Altogether, this work provides a unified, analytical framework for understanding BTSP in relation to network-level learning, establishing a theoretical foundation through which we can explore how this unique form of plasticity can be integrated into hippocampus-mediated learning processes.

## Results

### Generalized BTSP Rule Recapitulates Experimentally Observed Plasticity Kernels

We begin by building a generalized learning rule, based on experimental observations of BTSP. For clarity, we will hereafter refer to our rule as “gBTSP” (generalized BTSP, **Figure 1a-c**) and refer to the experimental phenomena as simply “BTSP”. To start, consider the simple case of a single postsynaptic neuron which triggers an instantaneous plateau event, and a single presynaptic neuron which fires a spike (**Figure 1a**). We assume the postsynaptic plateau updates weight *W* via some function, *W*_*kernel*_, which depends on the timing of the presynaptic spike relative to the plateau, i.e. Δ*W* ∝ *W*_*kernel*_(*t*_*pre*_ − *t*_*plateau*_).

**Figure 1.**
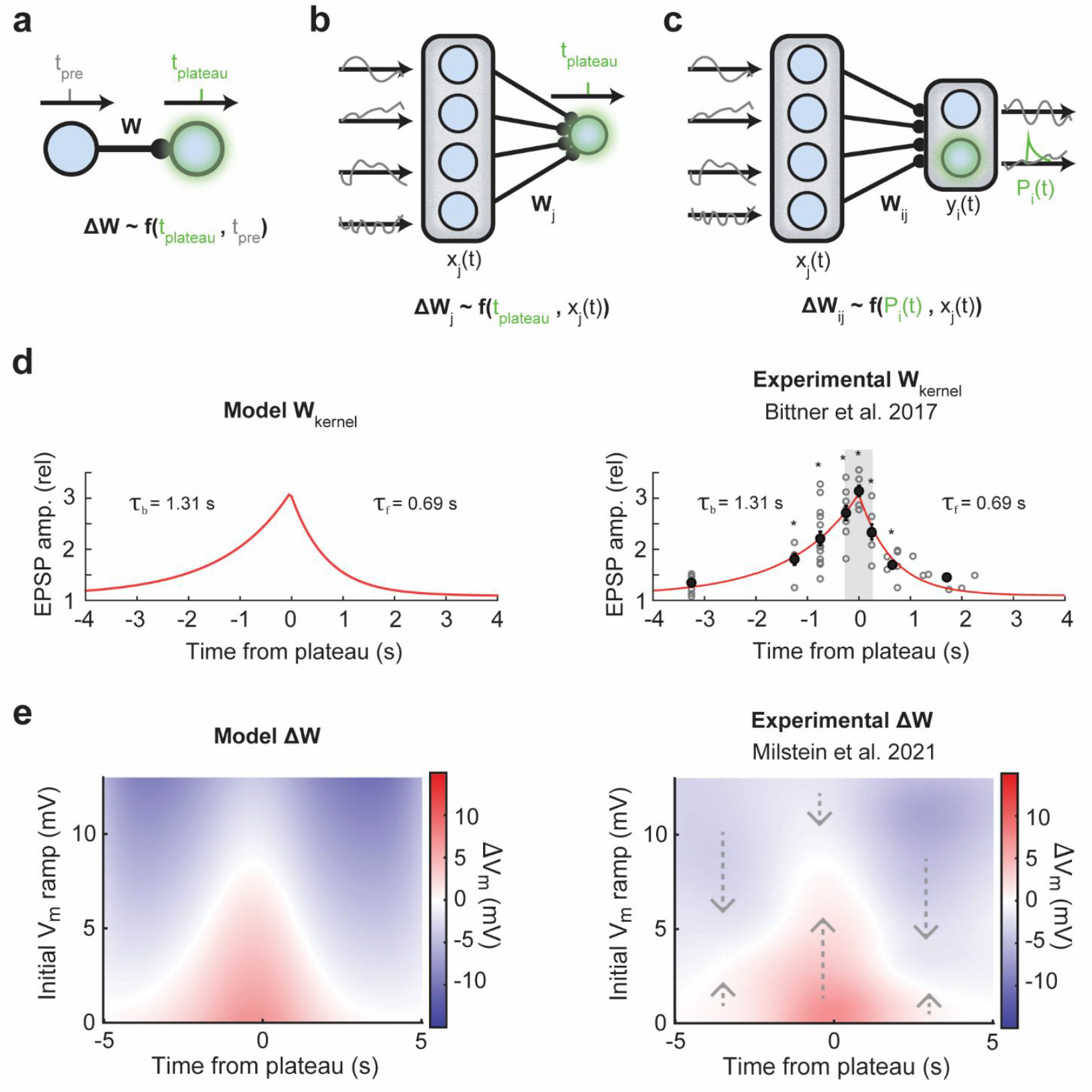
Generalized BTSP recovers experimentally observed plasticity kernels. **(a-c)** Schematics of different BTSP induction setups. **a)** A single plateau occurs in a postsynaptic neuron, and a single spike occurs in presynaptic neuron. Weight changes Δ*W* depend on some function of the relative time between the plateau and the presynaptic spike. **b)** A single plateau occurs in a postsynaptic neuron at *t*_*plateau*_, but now presynaptic neurons have some activity *x*_*j*_(*t*). Weight changes Δ*W* can be described as some function of the weight kernel and presynaptic activity. **c)** Potentially continuous plateau activity *P*_*i*_(*t*) occurs in the postsynaptic population. The resulting plasticity (“generalized BTSP” or “gBTSP”) depends on the weight kernel, presynaptic activity, and the postsynaptic plateau activity. **d)** Left, the kernel in our model uses two decaying exponentials, each with a different time constant. Right, the experimentally observed kernel, reprinted with permission from Bittner et al. 2017^2^; copyright AAAS. **e)** Left, the Δ*W* in our model when using the weight kernel from panel d) and a set of place fields (putatively from CA3) as presynaptic inputs. Right, observed Δ*W* in vivo, reprinted with permission from Milstein et al. 2021^1^; CC BY 4.0.

Owing to the wide temporal window in which plateau potentials have been observed to potentiate and depress inputs^1–3^, we assume that *W*_*kernel*_ operates on a timescale much larger than that of mere pre-post activity correlations. Specifically, we choose a *W*_*kernel*_ such that the application of our learning rule matches observed plasticity following application of a single plateau and bursting inputs in vitro (**Figure 1d**)^2^. Next, we relax our previous assumption that there is a single presynaptic neuron which fires a single spike, instead considering continuous presynaptic activity (of each unit *j*), *x*_*j*_(*t*) (**Figure 1b**). Now, the change in weights following a single plateau is a function of both the weight kernel and the presynaptic activity, i.e. Δ*W*_*j*_ ∝ *f*(*W*_*kernel*_(*t* − *t*_*plateau*_), *x*_*j*_(*t*)). See Methods for full derivation and expression.

To match experimental data showing that the amplitude of the formed field depends on the initial membrane voltage of the postsynaptic cell^1^, we also add in a dependence on the synaptic strength prior to the plateau event (see Methods). Following these additions to our rule, we can now use the same *W*_*kernel*_ from **Figure 1d** and show that for place field-like inputs *x*_*j*_(*t*), our rule recapitulates the observed plasticity kernels measure from single plateaus in vivo (**Figure 1e**)^1^. With this framework, the previously observed asymmetric offset of observed plasticity (**Figure 1e**)^1^ is a direct consequence of the shape of *W*_*kernel*_ **Figure 1d** (see Methods).

Finally, we want to consider the case for which there are multiple postsynaptic neurons, each of which may have a plateau (or potentially multiple). So, we introduce *P*_*i*_(*t*), a function representing the post-synaptic plateau potential at time *t* for neuron *i*. Critically, this post-synaptic plateau *P*_*i*_(*t*) is used only for learning and is distinct from the post-synaptic network activity, *y*_*i*_(*t*). Now, the change in the weights will depend on the weight kernel, the presynaptic activity, synaptic strength, and post-synaptic plateaus, i.e. Δ*W*_*ij*_ ∝ *f*(*W*_*kernel*_(*t* − *t*_*plateau*_), *x*_*j*_(*t*), *W*_*ij*_, *P*_*i*_(*t*)) (**Figure 1c**).

The full form of this dependence is given by the following equation, which we will hereafter refer to as “generalized” BTSP, since it is derived from various steps of generalization from our initial fundamental assumption (Δ*W* ∝ *W*_*kernel*_(*t*_*pre*_ − *t*_*plateau*_)):

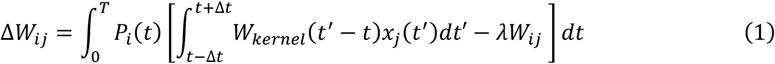

See Methods for full details and derivation. From this gBTSP equation, we can now take any set of inputs, choose any kernel, apply any arbitrary distribution of plateaus, and obtain a resulting change in the network’s synaptic weights. Note that gBTSP (as with BTSP) does not depend on postsynaptic activity directly, distinguishing it from standard Hebbian and Hebbian-like (e.g. STDP) learning rules.

Given this mathematical description of our learning rule, we now seek to gain a deeper understanding of its function. How does it operate inside of a network? What types of learning tasks is it well or poorly suited for? We will now investigate the properties of gBTSP in both unsupervised and supervised contexts, for both feed-forward and recurrent networks.

### Unsupervised gBTSP leads to competitive learning and one-shot field formation in feed-forward networks

To understand how gBTSP operates in the simplest case, we first consider gBTSP as an unsupervised rule. Returning to the simple case where we have a single plateau (and a short temporal kernel), Equation 1 simplifies greatly (see Methods), giving us an expression for plasticity of the form Δ*W*_*j*_ = *x*_*j*_(*t*_*plateau*_) − *λW*_*j*_. So, in this approximation, upon each plateau event, the weights would move towards a fixed point 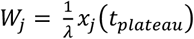. Such a formulation is reminiscent of classical conceptions of “competitive learning”^17–20^, whereby postsynaptic neurons *y*_*i*_ “compete” with each other to encode a pattern 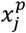 in its weights *W*_*ij*_ (where *p* is the index of a particular pattern, and *j* indexes over the pattern’s components). The decay term (− *λW*_*j*_) acts as heterosynaptic depression, promoting competition between units^21^.

Often, competitive learning is concerned with encoding *multiple* input patterns (or indeed, a whole distribution of possible input patterns), using various forms of “competition” (via some rule) to assign different postsynaptic neurons to represent distinct parts of the input space^17–20^. This algorithm has appealing aspects in the context of BTSP (and the hippocampus) – when presented with an input distribution, competitive learning can quickly (few-shot for a single unit) assign a unit to represent a part of that input space. Over the course of sampling the input distribution, a3 population-level representation slowly emerges. We might consider the hippocampus to be solving an analogous problem, e.g. forming a latent representation which tiles a given input space, taking care to have both a) coverage over the whole space, and b) well-separated or orthogonal latents which do not interfere with each other.

In order to adapt gBTSP for competitive learning, we must select a criterion for triggering plasticity events. Commonly, competitive learning methods only apply the weight update to the “best matching unit” (e.g. one which has a small Euclidean distance between 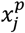 and *W*_*j*_)^19,22^. If we followed that logic, we would only trigger plateau events for these “best matching units”. However, this does not easily map onto learning in continuous time, particularly when we consider our temporally extended weight kernel. Instead, we choose an even simpler criterion, whereby a plateau event occurs in random neuron if the sum of total postsynaptic network activity *∑*_*i*_ *y*_*i*_(*t*) falls below some threshold *θ* (see Methods). In other words, if *∑*_*i*_ *y*_*i*_(*t*) < *θ*, we consider the currently arriving input **x(t)** to be poorly represented in the output layer **y(t)**. To amend this, the network fires a plateau, forming a new field (or translocating an existing one) that is tuned to **x(t)**. To test this simple algorithm for plateau assignment, we imagine a network of CA1 neurons to be receiving noisy but spatially tuned input from CA3 neurons, as an agent traverses an environment (**Figure 2a**). We consider both 1D (modelling an animal on a treadmill) and 2D (modelling an animal freely moving in a box) environments.

**Figure 2.**
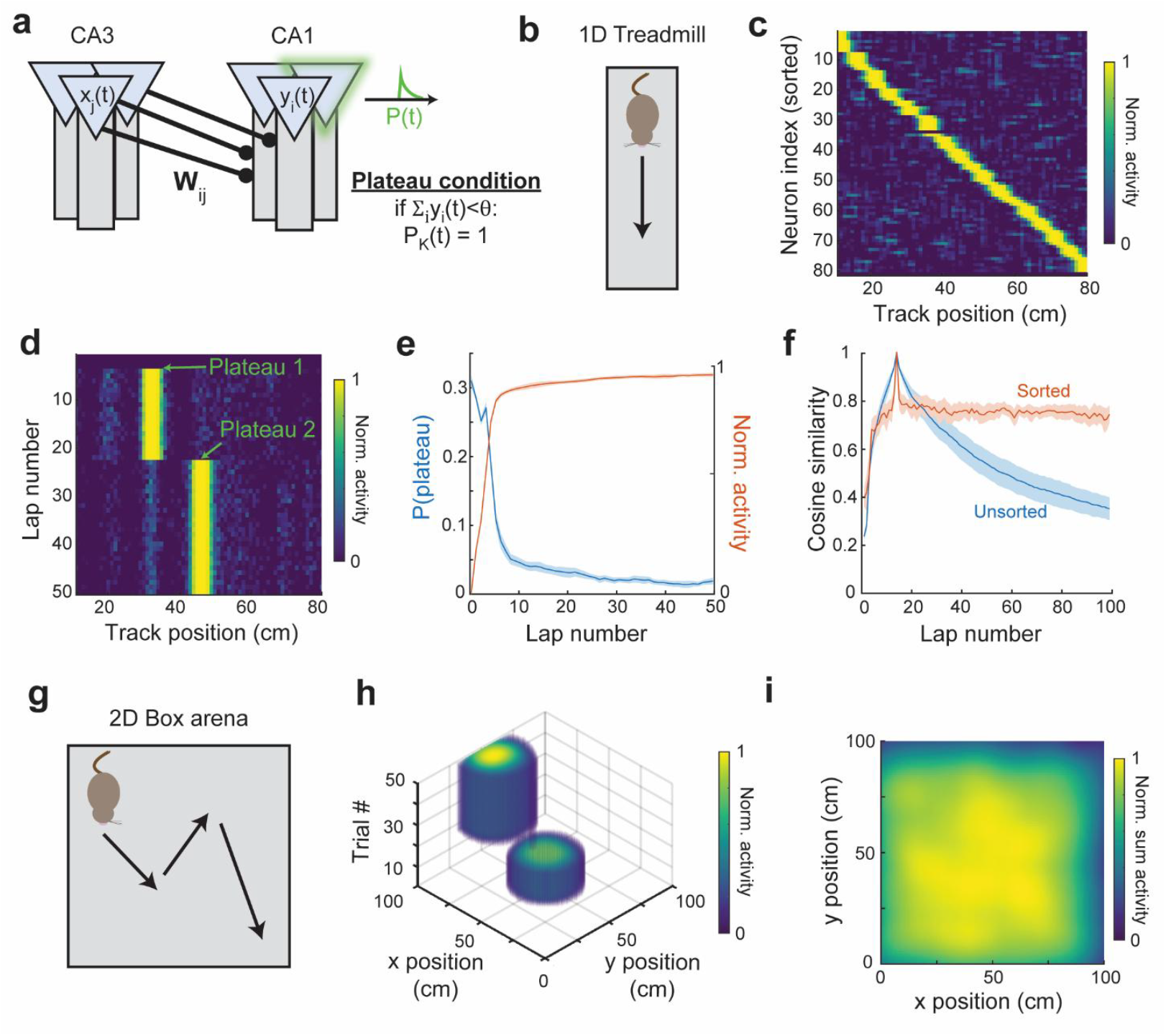
Competitive learning via gBTSP allows for one-shot formation of fields which tile the input space. **a)** Modelled inputs from CA3 project to our model CA1 neurons through weights *W*_*ij*_. A plateau fires at a random postsynaptic neuron when the sum of postsynaptic activity is below some threshold *θ*. **b)** For this task setup, inputs are drawn from a simulated agent running along a 1D treadmill. **c)** The learned fields across the population uniformly tile the 1D space. **d)** Using this plateau condition, single cells develop and translocate fields in a one-shot manner. For this particular unit, two plateau events occurred during training. **e)** The probability of a plateau event (blue line) peaks upon introduction to the novel environment, before decreasing to a baseline rate once a sufficient map has been learned. This time course is inverse to the total network activity (orange line). **f)** The baseline rate of plateau events (due to noise) causes representational drift. Blue, cosine similarity between unsorted network activity at the current lap and unsorted network activity at lap 10. Orange, cosine similarity between sorted network activity at the current lap and sorted activity at lap 10. Both measures peak at lap 10 because the cosine similarity of the activity at lap 10 with itself is 1. **g)** For this task setup, inputs are drawn from a simulated agent randomly exploring a 2D box. **h)** In a 2D environment, single fields still develop and translocate rapidly. **i)** Sum of neural activity for a trial where the agent explores the entire 2D environment. The learned latent representation covers the extent of the box.

For the case of a 1D environment, our agent moves along a treadmill at a uniform velocity (**Figure 2b)**, receiving spatially tuned inputs (see Methods), and applying a gBTSP plateau every time the low activity condition (*∑*_*i*_ *y*_*i*_(*t*) < *θ*) is met. After training, the population activity has evolved to span the space of the inputs, forming place fields which tile the length of the track (**Figure 2c**). Following the evolution of a single neuron in the network reveals that plateaus can both form and translocate fields in a one-shot manner (**Figure 2d**). The full, population level representation takes ~10 laps to mature, during which time there is a high likelihood of plateau events (as the network fills in “blank” spaces in the representation). The probability of plateau events scales inversely with the activity of the network, as we would reasonably expect from our criterion for triggering plateaus (**Figure 2e**). For subsequent laps after the network has evolved a mature representation (around lap 10), noise can still cause our plateau condition to be triggered. This can cause the translocation of existing fields (**Figure 2d**), also opening representational gaps that were previously filled. This effect leads to representational drift in unsorted representations, whereby the cosine similarity between the unsorted representation of the current lap and that of a reference lap (lap 10) increases as a function of experience (**Figure 2f, blue**), as has been reported experimentally^23,24^. However, this does not mean the content of the representation is fading – if we instead calculate the cosine similarity between sorted representation of the current lap and the sorted representation of a reference lap (lap 10), this measure remains stable over experience (**Figure 2f, orange**). This reveals that most of the representational drift occurring in the network is index-related, i.e. neurons may shift their tuning (or “label”) and “shuffle” where they occur in the sequence, but the internal, population-level sequential structure is maintained (**Figure 2c**)^25^.

We can extend further to a 2D environment (**Figure 2g**), where an agent takes a random walk inside a box, again receiving spatially tuned but noisy inputs (see Methods). Unlike the case of the 1D treadmill, where each lap the animal encountered the exact same input, here, the animal’s random walk means that it will experience a unique sequence of inputs each trial. Over the course of training, individual cells develop characteristic 2D place fields which evenly tile the space. Place field emergence in single-cells is still one- to few-shot, even in the 2D case (**Figure 2h**). We make the agent traverse the entire environment after training and find that the sum of network activity provides a map which covers the extent of the box environment (**Figure 2i**).

In summary, we find that for unsupervised learning in feed-forward networks, our mathematical formulation of BTSP can be mapped onto the classical framework of competitive learning. By applying gBTSP in simulated environments, we find that our network acts as we would expect from a competitive learner, taking a high-dimensional input space and summarizing it with a discrete set of lower-dimensional latent states. If BTSP indeed follows a simple threshold principle for competition, our model would predict that plateau probability across a network should be inversely proportional to that network’s activity (**Figure 2e**). Further, our model predicts that representational drift for a given learned neural trajectory is mostly a consequence of a “musical chairs-like” resorting, whereby transient and stochastic dips in total network activity in one location are likely to trigger a translocating plateau event, leading to a dip in network activity in the translocated field’s previous location (**Figure 2d-f**).

### Unsupervised gBTSP can facilitate attractor learning in recurrent networks

Given that BTSP has been observed in CA3, driving plasticity in recurrent CA3→CA3 synapses^4^, we now consider how unsupervised gBTSP might be understood in the context of a recurrent network. A common model of CA3 is that of an attractor network^26–30^, where an “attractor” can be framed in the context of discrete memory states (e.g. a Hopfield model)^31,32^, or a continuous manifold^27,33–35^. Experimental results have demonstrated CA3’s ability to both pattern complete and tune its activity via velocity-dependent inputs^4,36,37^, hallmarks of a (recurrent) attractor network. As such, it is reasonable to suspect BTSP may be involved with the formation or maintenance of these networks. Indeed, previous theoretical work has shown that the BTSP rule’s characteristic kernel is well suited for optimal memory storage in discrete memory networks^4^, but it remains unclear if/how an unsupervised form of BTSP can give rise to attractors.

In order to simplify our problem, we will utilize a two-part architecture in our network, inspired by similar parametrizations of recurrent nets designed to learn or sustain attractors^38–41^. In short, we imagine there to be two distinct populations in CA3, with only one of the populations eligible to receive plateaus via gBTSP. This assumption is based on experimental results which have shown that the ability or propensity of pyramidal cells in CA3 to have complex bursting events (i.e. a plateau) is variable from cell to cell, and may depend on features such as topographic position and/or dendritic morphology^42,43^. Both of our populations (“visible” neurons *u*_*j*_(*t*) and “seed” neurons *s*_*i*_(*t*)) connect recurrently to each other, via “encoding” weights **W**^**e**^ and “decoding” weights **W**^**d**^, with the visible neurons receiving external input *o*_*k*_(*t*), and only the seed neurons are eligible for plateaus (**Figure 3a**). The relative strength of recurrent/external input onto the visible neurons is governed by a gating function which depends on the norm of the external input (see Methods): when external input is high, recurrent input is low, and vice versa. Owing to this gating, high external input effectively turns the network into a feed-forward one, and when input is removed, the network restores its recurrency. This dual nature of the network allows us to take advantage of recurrent computation in the low-input phase, while making use of unsupervised, feed-forward gBTSP in the high-input phase. For full details, see Methods.

**Figure 3.**
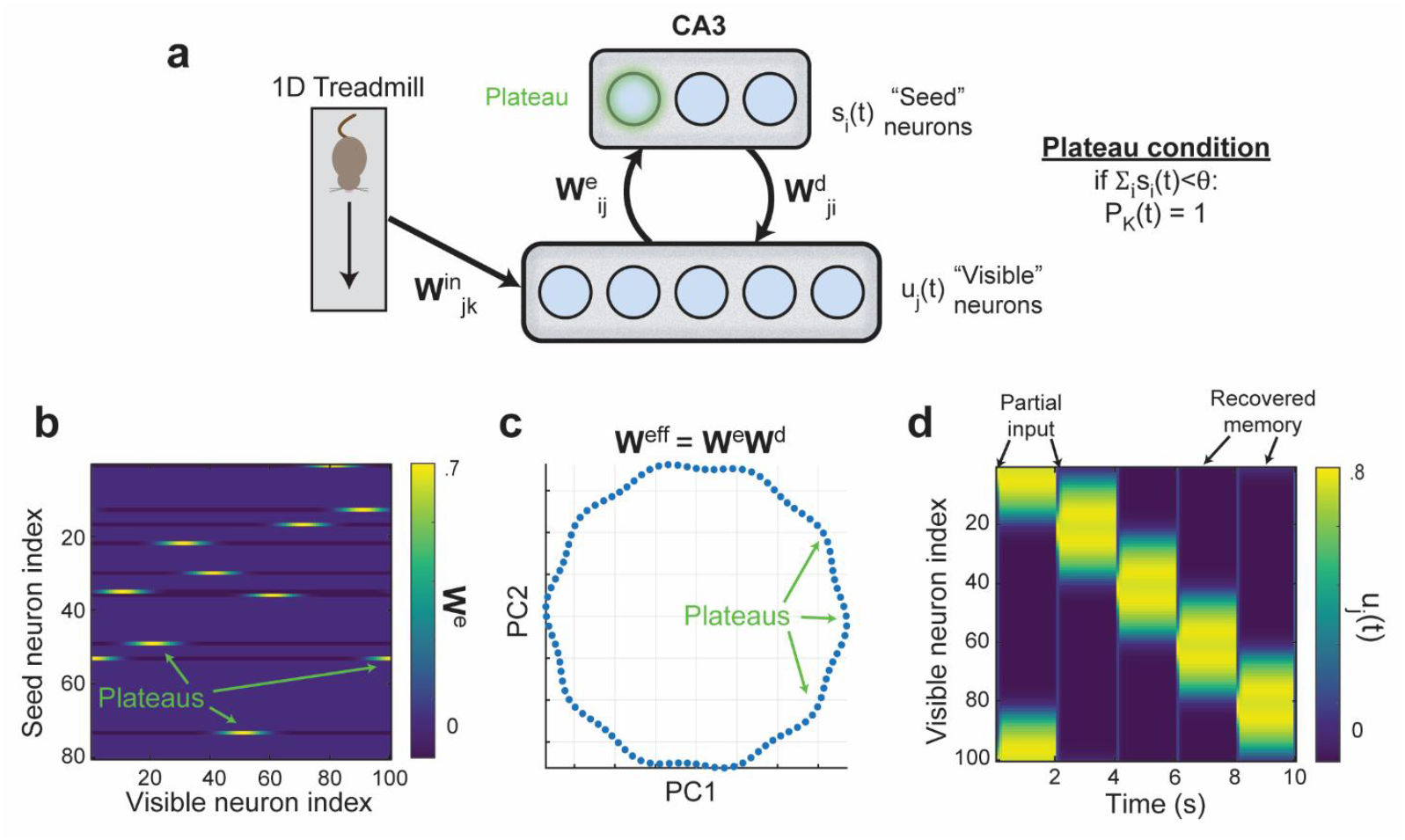
Building an attractor network via gBTSP. **a)** A set of spatially tuned noisy inputs are drawn from a simulated agent running along a 1D treadmill. These inputs project to “visible” neurons through a set of input weights. Visible neurons project to “seed” neurons via encoding weights, and seed neurons project back to visible neurons through decoding weights. There are no recurrent weights within each layer. This produces an effective recurrent weight matrix **W**^**rec**^ = **W**^**e**^**W**^**d**^. A plateau fires at a random seed neuron when the sum of postsynaptic activity is below some threshold *θ*. **b)** Plateau events create seed neurons which are sensitive to certain combinations of visible neurons. **c)** The effective recurrent weight matrix, **W**^**rec**^, forms a ring attractor, as viewed in the first two principal components. Each of the nodes along the attractor is a fixed point created by a plateau event. **d)** Partial inputs given to the network for a single timestep at times 0,2,4,6, and 8 seconds recover a memory state of the network, which persists until the next partial input is presented.

Since our aim is to learn attractor states (i.e. **u(t)** = **u(t** − **1)** for no input), we would like our effective recurrence, **W**^**rec**^ = **W**^**d**^**W**^**e**^ to be approximately to the identity matrix. To avoid trivial solutions, we make two choices. First, we set the seed population to be smaller (in number) than the visible population, forcing the network to compress and then decompress its representations (this is equivalent to making **W**^**rec**^ low-rank). Second, we apply the same competitive learning framework from the feed-forward case (*∑*_*i*_ *s*_*i*_(*t*) < *θ*) to learn the encoding weights, so that the seed neurons learn latent representations which tile the input space. The decoder weights are set to be the transpose of the encoder weights, which is sufficient since the learned encoding is well-separated (near orthogonal). Ideally, some biophysically plausible learning rule can govern the evolution of these decoder weights^44^, but for the purposes of this study, we use the transpose relationship as a simple approximation.

We simulate an agent running along a 1D treadmill, again receiving spatially selective inputs which are processed by our model CA3 network (**Figure 3a**). The seed neurons are allowed to plateau, doing so under the same low-activity criteria as in the feed-forward case. During training, the agent runs along the track and external inputs are strong. Plateau events occur in response, guiding the evolution of the encoding weights (and thereby the decoder weights). Seed neurons form receptive fields to their visible neuron counterparts (**Figure 3b**), similar to the formation of place receptive fields in the purely feedforward case (**Figure 2c**). After just the first lap of training, the recurrent weights have formed a ring topology, with fixed point nodes (memories) at locations dictated by the plateau events (**Figure 3c**). This topology is well explained by the first two principal components (**Supplemental Figure 1**). So long as external input continues, the network remains largely feed-forward, but upon removal of external inputs, the network is dominated by recurrency, and relaxes into one of its learned fixed points. If partial and intermittent inputs are given, the network can switch between these encoding (feed-forward) and recall (recurrent) modes repeatedly, recovering a new memory each time it samples its inputs (**Figure 3d**).

Altogether, by using a low-rank formulation of our network recurrency, and gating plateau events to occur in a certain subpopulation of our network, we demonstrated that gBTSP can play a crucial role in the rapid formation of an attractor network. Such a role would be consistent with observations of BTSP in CA3^10^, a region often hypothesized to play the role of an attractor^26–30^. Future experimental and theoretical work can further illuminate the functional structure of CA3 and the role BTSP plays in forming and maintaining attractor states.

### Supervised gBTSP can support rapid task learning in feed-forward networks

While competitive learning provides an unsupervised framework by which we might understand the function of BTSP during novel, unguided exploration, it still leaves unanswered what role direct supervisory credit assignment might play. A popular hypothesis for BTSP posits that EC3 inputs to the distal dendrites act as supervisory “targets”, which in turn trigger plateau events so that the somatic activity can match this dendritic target^5^. However, it is not clear what these targets are, i.e. “when” and “where” should a plateau event occur? To rephrase the question in a more quantifiable way: if we define a given loss ℒ as the mean squared error between the network output *y*_*i*_(*t*) and some target *ŷ* _*i*_(*t*), when and where should we trigger plateau events to minimize this loss? Equipped with our learning rule, we have the tools to answer this question. To this end, we set our expression for Δ*W*_*ij*_ from gBTSP (Equation 1) to be equal to expressions for Δ*W*_*ij*_ from traditional supervised learning (in the simplest feed-forward case, the “delta rule”), and solve for *P*_*i*_(*t*) (see Methods). In the case of a simple feed-forward network, that expression is the following:

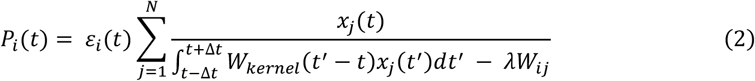

Where is our *ε*_*i*_(*t*) = *ŷ*_*i*_(*t*) − *y*_*i*_(*t*) task error. This expression gives the plateau function *P*_*i*_(*t*) which will descend the loss gradient on a given trial. Equipped with this formula, we can now test the ability of gBTSP to learn in supervised learning contexts.

As a sanity check, we first consider the trivial case where our inputs are already spatially selective (such as those arriving from CA3)^1,14^ and our output represents a single CA1 pyramidal cell subject to gBTSP (**Figure 4a**). We choose a target function *ŷ* (*t*) that is a putative place field, modeled as a Gaussian bump centered at a specific location in the environment (see Methods). We find that the network can match this target through plateau-driven learning (**Figure 4b**), demonstrating the fundamental capability of gBTSP to adapt network weights toward a desired output function (**Supplemental Figure 2**). The plateau function which solves the task is, as expected, centered at the location of our target function, and is most significant within the first 3-5 trials. The field itself also rapidly emerges on this same timescale (**Figure 4c**), in agreement with experimental results where place fields were formed via the artificial induction of plateaus a) at the location of the desired place field, and over only a few (<10) trials^1–3^.

**Figure 4.**
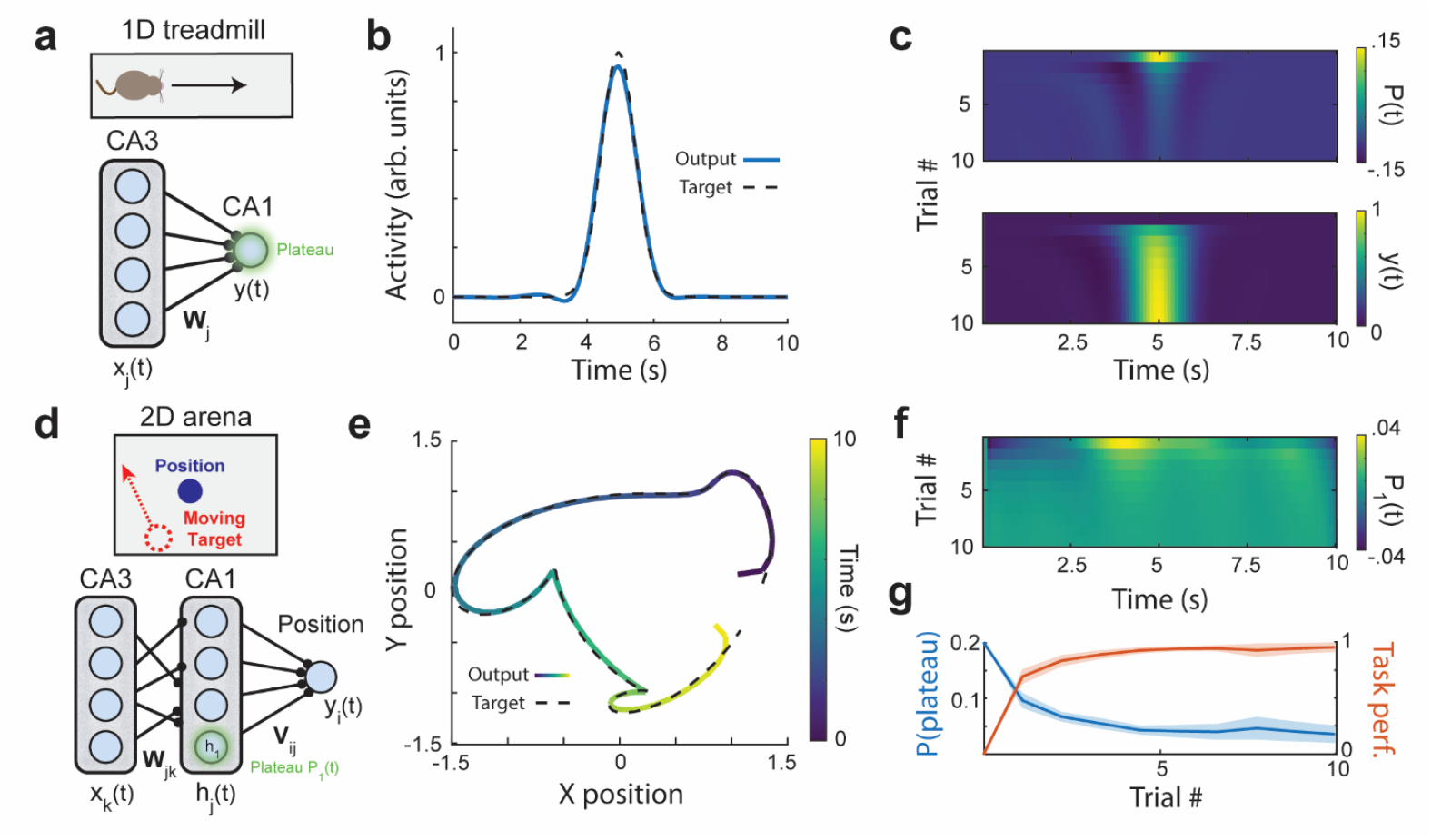
Feed-forward network rapidly acquires task via supervised gBTSP. **a)** Feed-forward network with inputs *x*_*j*_(*t*) project to output *y*(*t*) via weights *W*_*j*_. The inputs *x*_*j*_(*t*) are spatially selective and represent an animal running along a 1D track. **b)** A unimodal gaussian target function (dotted black line) and the trained output (blue line) after the first 10 trials of gBTSP training. **c)** Top, the plateau function *P*(*t*) over the first 10 trials of training. Middle, the output *y*(*t*) over the first 10 trials of training. Bottom, the output activity *y*(*t*) over the first 10 trials of training. **d)** A two-layer network with inputs *x*_*k*_(*t*), hidden units *h*_*j*_(*t*), and 2D output *y*_*i*_(*t*) which represents location. The target trajectory moves in a set 2D path each trial, and the network must learn to track the target. **e)** The target path (dotted black line), and the learned path (colored line). The color here represents the time at which the agent is in a given location. **f)** Plateau function for the first unit in the hidden layer, *P*_1_(*t*), over the first 10 trials of training. **g)** Task performance (proportional to 1 - *ε*_*i*_(*t*)) and plateau probability over the first 10 trials of training.

Next, we test our ability to train the network on a more complicated task, in a network with a single hidden layer which is subject to gBTSP. In this task, we model an agent learning to match its location in a 2D arena to some target trajectory in that arena (indicated, say, by targeted illumination). One can consider this task as a navigation-based analogue to smooth pursuit or continuous reaching tasks. Rather than merely generating static spatial patterns, the network now must take a dynamic input *x*(*t*) and learn to generate a dynamic 2D position output *y*(*t*) (see Methods, **Figure 4d**). After the first 10 trials, the agent has learned to track the target trajectory (**Figure 4e**). Unlike the previous example, where the plateau location was obvious by design, here it is unclear a priori when and where plateaus should occur in the hidden layer to solve the task. We find that a more complex pattern emerges for the plateau function in a sample neuron, and there is no longer a simple correlation between the network target and the shape of its plateaus (**Figure 4f**). This is because our expression for the plateau function (Equation 2) will depend on the backpropagated error (see Methods) in networks with more than one layer. Finally, our model predicts that, in a supervised framework, the probability of plateau events in the full population should be inversely correlated with task performance (or positively correlated with task error), decreasing over the course of task learning (**Figure 4g**).

Together, these results demonstrate that we can use gBTSP to descend the gradient of a supervised loss. In other words, the algorithm “distributes” plateau events to certain neurons at certain times in order to optimize overall network performance. If EC3 does indeed dictate plateau induction via supervised “targets”, as has been suggested^5^, then our framework provides a computational tool to understand and potentially infer the content of these signals. One testable prediction our supervised framework makes is that plateau probability should rise and fall with the inverse of task performance (**Figure 4g**).

### Supervised gBTSP in recurrent networks fails to support rapid, one-shot learning

Finally, we wish to examine the feasibility of our rule in a fully recurrent network (to mimic CA3), learning a supervised task which requires maintenance of an internal memory (via recurrency). Our network consists of hidden units with activation *h*_*j*_(*t*), recurrently connected via weights 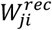, and projected to output *y*(*t*) via weights 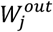 (**Figure 5a**). Internal weights 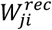 are trained indirectly through plateau induction, which is dictated by a recurrent update rule, that we derive by comparing our gBTSP weight update to that of backpropagation through time (BPTT) (see Methods).

**Figure 5.**
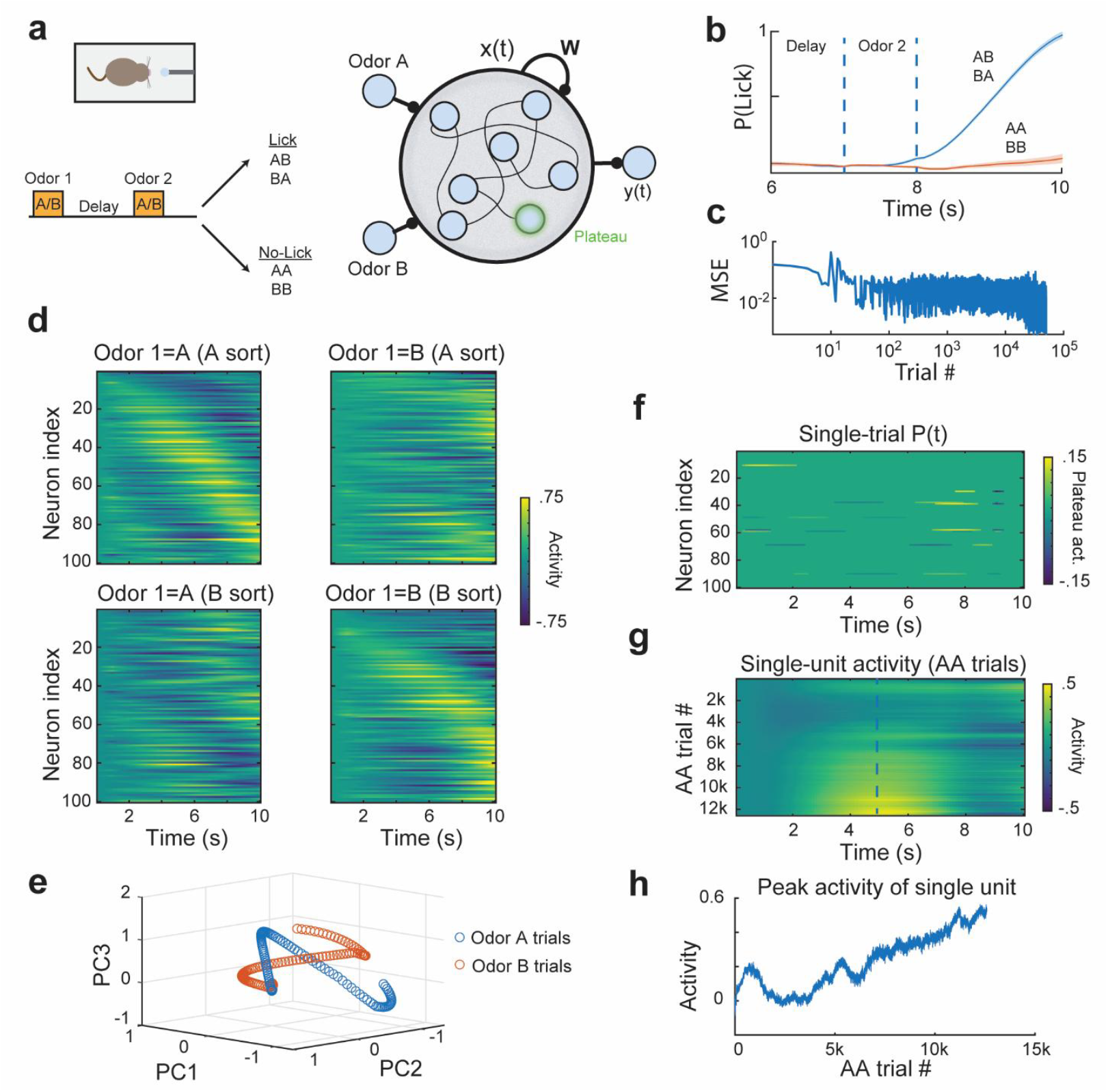
Learning a complex recurrent task with gBTSP requires slow and precise credit assignment. **a)** Simulated agents are trained on a delayed-non-match-to-sample (DNMS) task where they must distinguish between sequential “odor” pairs. The agent must learn to “lick” for non-matching sequences (AB, BA), and refrain from licking for matching sequences (AA,BB). Right, the model consists of a recurrent network with two odor inputs, hidden activities **x(t)** and recurrent weights **W**. The hidden units project to an output **y(t)** via weights **V**. Recurrent weights **W** are trained via plateaus occurring in the hidden units according to our gBTSP algorithm (see Methods). Output weights are trained via the delta rule. **b)** After training, the model output has learned to lick following odor 2 in the AB/BA trials while refraining from licking in the AA/BB trials. **c)** Mean squared error (MSE) decreases over training, demonstrating successful learning, albeit across tens of thousands of trials. **d)** Neural activity patterns across all 100 neurons, averaged over trials which began with odor A (left column), or odor B (right column). The neurons are sorted by time of maximum activity in the odor A trials (top row), or time of maximum activity in odor B trials (bottom row). The activity maps reveal the network has learned distinct sequences of activity for the initial odors, thereby forming a working memory of the first odor’s identity. **e)** Activity of each trial type (odor A or odor B), projected onto the first 3 principal components of the total activity space after training. For variance explained, see (**Supplemental Figure 3**). **f)** Single-trial plateau events across neurons and time, showing sparse activation. **g)** Evolution of activity for a representative single unit during AA trials over training. The final neural tuning slowly develops over many thousands of trials. **h)** Activity of the same representative unit at t = 5 seconds, across AA trials, again highlighting gradual field development.

We illustrate the behavior of the model using a standard delayed-non-match-to-sample (DNMS) task, where our simulated agent must distinguish between sequential pairs of odors, and only “lick” in responses to non-matching pairs (AB, BA), refraining from licking following matching pairs (AA, BB) (**Figure 5a**). We choose this task because it requires the network to maintain a memory of the first odor’s identity (by leveraging recurrent learning). Moreover, previous experimental work has shown that animals trained on the same task developed two distinct hippocampal sequences of activity which encoded the identity of the first odor^45^.

To avoid instabilities during training, we combine our update with an adaptive optimizer (ADAM) before updating the weights of the network (see Methods)^46^. We find that gBTSP can learn the target function, choosing to “lick” when the two samples are non-matching, and forgoing licking when the two samples match (**Figure 5b**). However, unlike the simpler tasks we have thus far described, training a recurrent network on the DNMS task takes many thousands of trials (**Figure 5c**). In order to solve the task, the network develops distinct internal representations for the cases when Odor 1 = A, and when Odor 1 = B (**Figure 5d**). These representations are well explained by their first three principal components, with Odor 1 = A trials and Odor 2 = B trials making distinct trajectories in this subspace (**Figure 5e, Supplemental Figure 3**). These distinct representations act as a memory trace of the first odor’s identity, thereby allowing the network to judge “match” vs. “no-match” upon presentation of the second odor. These representations resemble neural sequences observed in experimental studies^45^, but as in previous theoretical work^47^, we found that adding a ramping component to the task target best recovered this sequential activity (see Methods). The plateaus in the network which facilitate learning were constrained to be stochastic and sparse, with only 10% of neurons allowed to plateau on a given trial, and only events which crossed an absolute magnitude threshold contributing to learning (**Figure 5f**). Though these constraints may result in single trial samples of P(t) which share the sparse nature of plateaus observed in vivo^5,48^, when we examine the evolution of single cell fields, we see that they develop very slowly, taking thousands of trials (**Figure 5g,h**). Another alternative would be to increase learning rates, but doing so results in unstable learning (**Supplemental Figure 4**). In short, spatiotemporal credit assignment in recurrent networks is notoriously difficult^49^, and unsurprisingly, solving for *P*_*i*_(*t*) via gBTSP (as opposed to solving for *W*_*ij*_ directly via BPTT) does not bypass these limitations. In other words, the standard tool of gradient descent does not stably resolve the question of “where” and “when” BTSP events should occur in a recurrent network in order to solve a supervised task. We will now elaborate on a more complete answer and discuss how we can reconcile these apparent hard limits on the speed of learning with the existence of BTSP in CA3, a highly recurrent network.

### Rapid activity changes due to gBTSP are fundamentally limited in deep and/or recurrent networks

In shallow feed-forward networks, gBTSP could recover few-shot learning as observed experimentally with BTSP. However, as we have demonstrated particularly for supervised learning in the recurrent network, single-cell learning via gBTSP was very slow. Moreover, attempting to speed-up learning results in instabilities (**Supplemental Figure 4**). Why is this?

For the following, we will step aside from the specifics of gBTSP to make a more general formulation of the problem. Let us assume only that a) plateau events exist, and b), they cause single-cell activity to change by a fixed amount Δ*x*, remaining agnostic about the type(s) of learning involved in bringing about this change Δ*x*. We can consider learning via these plateau events from the perspective of optimizing within a loss landscape, taking a step Δ*x* along the direction of the descending gradient (first derivative). In landscapes with “fine-grained” or “sharp” features, a step of size Δ*x* can overshoot the global minimum (**Figure 6a**). Conventional learning approaches address this issue by reducing step sizes (i.e. taking a step of size δ*x* < Δ*x*) (**Figure 6ai**), thereby allowing learning to converge to the minimum.

**Figure 6.**
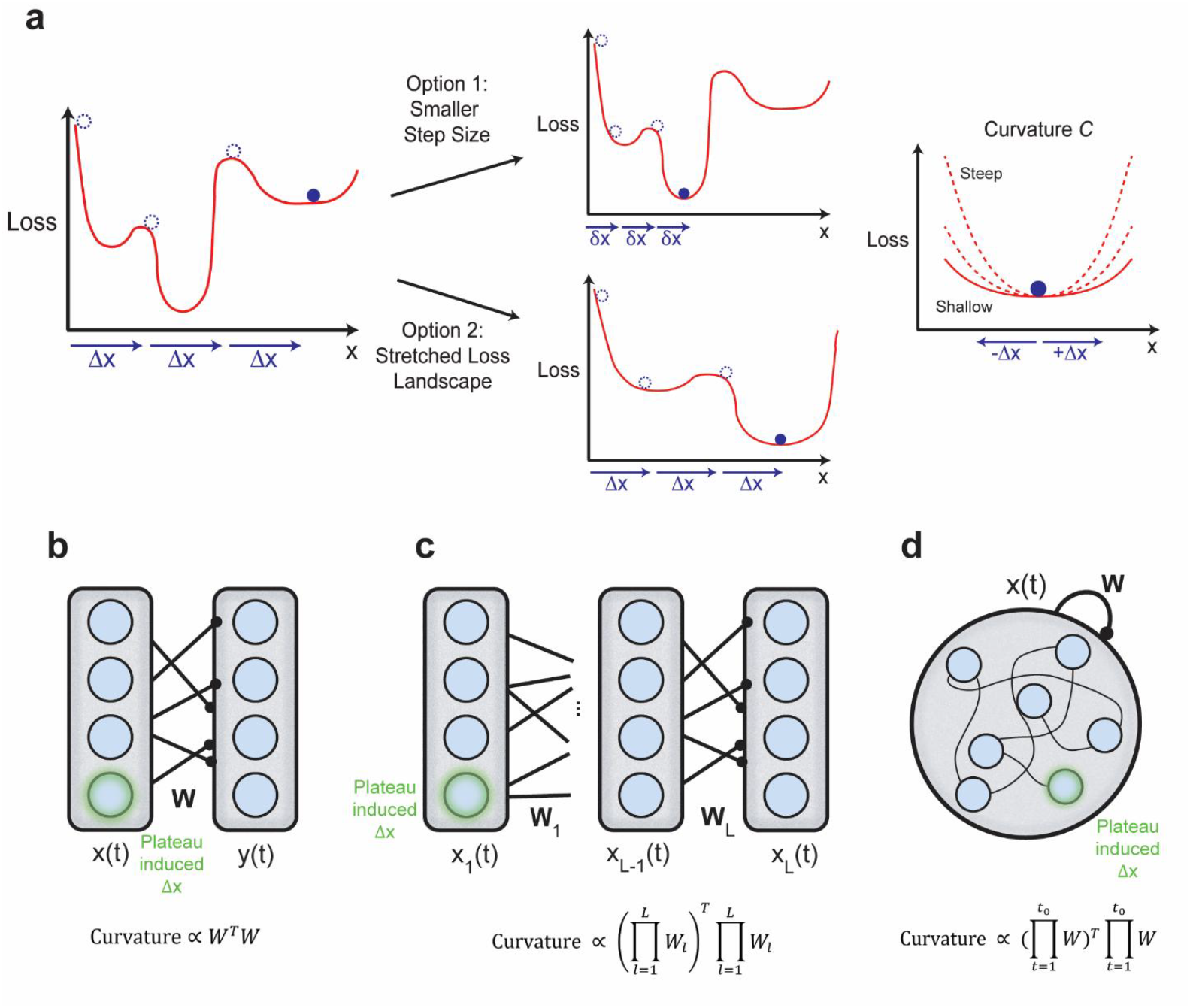
Shallow losses are required for few-shot learning. **a)** Red, an arbitrary loss function ℒ(*x*) which depends on the network state, *x*. In order to learn, the network is restricted to make discrete jumps of size Δ*x*. Blue dotted circles, previous values of ℒ(*x*). Filled blue circle, current value of ℒ(*x*). Top middle, smaller discrete jumps of size δ*x* are sufficient to reach our learning objective, but results in a slow evolution of network activity over learning (“dough” learning, right). Bottom middle, stretching the loss function also allows us to reach our learning objective, while maintaining a fixed step size Δ*x*. Single unit activities rapidly change over the course of learning, even if the population representation evolves slowly (“popcorn” learning). This stretching picture requires a shallow but non-zero curvature (right). **b)** Shallow feed-forward network, and its associated curvature. **c)** Deep feed-forward network, with L layers, and its associated curvature. **d)** Recurrent network which runs for T timesteps, and its associated curvature. Note that this picture can be related to that of the deep feed-forward network, if we imagine each “layer” to be the activity of the recurrent net at a time t, and the weights between these layers to be the shared weight matrix W.

An alternative approach involves modifying the loss landscape itself. By “stretching” the landscape, the same step size Δ*x* becomes proportionally smaller relative to the landscape features, “smoothing” out sharp features preventing overshooting (**Figure 6aii**). Mathematically, this “stretching” operation locally shrinks both the first and second derivatives of the loss with respect to activity. Since we are assuming each optimization step takes a fixed step size Δ*x* regardless of the gradient (first derivative) amplitude, we can focus on conditions on the second derivative (which we will hereafter refer to, for simplicity, as the local “curvature” *C*), and show how this curvature depends on features of the network. Intuitively, learning in this “stretched” landscape might be considered akin to the evolution of microwave popcorn, in the sense that while population-level representations (the popcorn bag) may evolve gradually, individual units (the kernels) undergo rapid, stochastic transitions to their final states on timescales significantly shorter than the overall system evolution. To support this “popcorn”-like approach (which we posit to be more BTSP-like), local curvature must be small (i.e. the loss must be locally “shallow”) to prevent overshooting, but non-zero to enable learning in the first place.

In the case of a single-layer feedforward network, inputs *x*_*j*_(*t*) project to output *y*_*i*_(*t*) = *∑*_*j*_ *W*_*ij*_*x*_*j*_(*t*). (**Figure 6b**). “Plateaus” of size Δ*x* occur at the inputs, and the loss is the mean squared error between output **y** and target *ŷ*. Solving for the curvature, we find that *C* ∝ *W*^*T*^*W* (see Methods). It is trivial enough to construct a network where *W*^*T*^*W* is small but non-zero. For example, if *y*_*i*_(*t*) receives many inputs, each with a small weight *W*_*ij*_, any change in a single input *x*_*j*_(*t*) will lead to small change in *y*_*i*_(*t*). So, in the case of a single layer feed-forward network, a small but non-zero curvature is achievable, meaning rapid changes in activity arising from BTSP can lead to stable learning.

If we consider a deep feed-forward network with layers *l* and layer specific weights *W*_*l*_ (**Figure 6c**), the expression for the curvature becomes more complicated, depending on products of all the layer-specific weights together in sequence 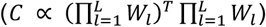. Unfortunately, it is not trivial to make these products small but non-zero - in fact, they are the same troublesome mathematical objects which lead to the problem of exploding and vanishing gradients in gradient descent^49–51^.

Recurrent networks (**Figure 6d**) can be conceptualized similarly to deep feed-forward networks (**Figure 6c**), but with each “layer” representing a different timestep in the network, with the weight matrix *W* is applied at each timestep. In turn, curvature of the loss in a recurrent network depends on similar products 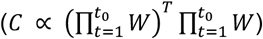 which also lead to exploding and vanishing contributions from an update Δ*x*.

These conditions set fundamental limits on both the architectures and the tasks for which deep or recurrent artificial networks can support rapid changes in single-unit activity. Despite these theoretical limitations, few-shot BTSP events have been observed in CA3^4^, which is highly recurrent. The analysis above is idealized, and biological neural networks may have yet unknown mechanisms which allow them to bypass these restrictions. However, if we hypothesize that recurrent connectivity in CA3 is indeed subject to these constraints, specialized architectural features would be required to stabilize learning dynamics and prevent vanishing or exploding effects arising from rapid plasticity events driven by BTSP.

## Discussion

The discovery of Behavioral Timescale Synaptic Plasticity (BTSP) unearthed an apparent paradox in our understanding of learning in the hippocampus. One the one hand, successful models of complex population-level hippocampal function (e.g. the formation of cognitive maps^52–54^) depend critically on recurrent computation, and in turn, seem to depend on the slow, precise training of recurrent weights. On the other hand, experiments in hippocampus observe a learning rule (BTSP) which is very fast and has a very distinct lack of temporal specificity. BTSP’s hallmark features—its wide temporal kernel spanning seconds and its rapid, one-shot field formation—run counter to conventional wisdom that precise, gradual weight changes are necessary for stable learning. To examine the computational implications of BTSP, we proposed a generalized mathematical framework, gBTSP, for which plasticity is governed by wide temporal kernels and a postsynaptic “plateau function” P(t). We test its properties across different network architectures (feed-forward and recurrent) and learning paradigms (supervised and unsupervised).

We demonstrated that unsupervised gBTSP in feed-forward networks maps well onto the framework of competitive learning, wherein neurons “compete” to represent distinct regions of the input space. This framework accounts for experimentally observed phenomena, including the rapid formation of place fields and their distribution across the environment. If we assume BTSP is operating according to unsupervised principles, our model predicts that plateau probability should inversely correlate with network activity, offering a testable hypothesis for future experiments. Moreover, we found that individual neurons can undergo rapid remapping while the population-level representation maintains coverage of the environment—exhibiting representational drift primarily through index “shuffling” rather than degradation of the underlying representation. However, we only consider a few hundred trials of unsupervised learning, so it remains unclear to what extent this model can explain recent experimental results regarding place field stability over days^55^ or over the course of goal-oriented spatial learning^56^. One could also imagine extending our framework by considering other forms of competitive learning, such as self-organizing maps^19,57^, which assume an underlying functional structure which is maintained during learning. Such an extension might explain the observations of functional clustering of field formation events around “seed” plateau neurons^58^, and other topographically related phenomenon observed in plateau generation^59^.

In recurrent networks like CA3, we demonstrated that unsupervised gBTSP can facilitate attractor learning when implemented with appropriate architectural constraints. This aligns with previous theoretical work showing BTSP’s suitability for memory storage in discrete attractor networks^4^. To preserve stability during learning, we used a low-rank parameterization of the network’s recurrent weights, only applying gBTSP to the “encoding” portion of this parameterization. However, there are also other promising avenues for considering BTSP in the context of unsupervised, recurrent learning. For example, under certain conditions STDP in a recurrent network can approximate Hidden Markov Model Learning, a very powerful tool for discovering underlying latent structure^60^. Recent experimental work which recorded hippocampal activity over the course of learning observed an orthogonalization of the latent map^61^ – a feature they found was best described by networks which learned HMMs^60,62^, including a version of the STDP-recurrent network. Although BTSP learns with a much larger temporal kernel than STDP, they have a similar fundamental structure. One can imagine that a mapping of BTSP onto HMM learning may be possible, though the rapid learning and large temporal kernel of BTSP present non-trivial challenges to stability and convergence.

For learning based on explicit error functions (i.e. supervised learning), we derived analytical expressions that determine when and where plateau events should occur to optimize task performance. This formulation allows us to understand BTSP in the context of gradient-based learning, with the plateau function effectively distributing “credit” for errors across the network. Further, because we have an explicit analytical expression for our plateau function, we can constrain it to be stochastic and sparse, akin to BTSP events observed in vivo^5,48^. We show that gBTSP can successfully learn feed-forward tasks, while retaining key features of observed BTSP, such as few-shot learning.

While our supervised gBTSP successfully learned complex tasks in feed-forward networks, maintaining the rapid learning characteristic of BTSP, deep recurrent networks proved more challenging. We showed that these challenges arise from fundamental stability limitations of large, rapid activity changes in deep and recurrent networks, limitations which are similar mathematically to exploding or vanishing gradients in backpropagation^49–51^. This presents an apparent paradox, as BTSP has been experimentally observed in the highly recurrent CA3 region. Perhaps structures such as so-called orthogonal or unitary networks, which preserve spectral norms of the recurrent weights (and thereby maintain stable gradient flow), can offer a solution, but training in these networks is difficult to reconcile with gBTSP^63–65^. Alternatively, it might be sensible to model CA3 as dynamically regulating its recurrence, gating certain pathways such that they behave as feed-forward networks during learning episodes (note that we used this method earlier to train our unsupervised recurrent network; **Figure 3**).

Our analysis has focused on a “strong” hypothesis, by which hippocampal learning is governed mainly via BTSP, and the plateau events follow some unifying principle (such as minimizing a particular error or loss function). We call this the “strong” assumption, because it remains unknown what fraction of overall learning is due to discrete, rapid BTSP events, it is highly likely that multiple forms of plasticity, including BTSP, are active simultaneously, and c) it is unclear if plateau-driven learning is guided by any sort of governing computational principle. A “softer” hypothesis might posit that BTSP is but a small fraction of hippocampal learning, and/or it is relegated to a trivial function such as taking mere random “snapshots” of complex representations occurring in other cortical areas. While this “soft” hypothesis remains worthy of consideration and further study, rates of BTSP appear to be relatively high, particularly in novel environments^5,48^, and previous theoretical work suggests that purely random plateau occurrence is inconsistent with the task-specific formation of complex hippocampal representations (i.e. splitter cells)^66^.

The limitations we identify suggest that while BTSP may indeed play a crucial role in hippocampal learning, at the very least, its implementation likely requires specialized circuit designs and/or other forms of plasticity to maintain stability. Still, the stark contrast between traditional gradient-based learning (where both population and single-unit representations evolve gradually), and BTSP-like learning, (where individual units can change rapidly while population representations evolve more gradually), highlights a fundamental difference between learning in artificial and biological systems.

Ultimately, this work introduces a generalized mathematical and analytical framework for BTSP (gBTSP) and uses this framework to investigate how plateau events may be distributed to solve learning tasks. Our findings suggest that while placing the entire burden of credit assignment on plateau events alone may be insufficient to explain complex aspects of hippocampal learning, BTSP is capable of rapid memory formation and latent encoding, particularly in feed-forward, and constrained recurrent networks. Future work should identify the biological circuits and plasticity mechanisms that stabilize hippocampal networks undergoing BTSP, particularly within CA3, to better understand how BTSP contributes to the development of hippocampal cognitive functions.

## Methods

All parameters for the following methods are included in **Table 1**.

**Table 1:** Model Parameters.

| Parameter | Value | Units | Description |
| --- | --- | --- | --- |
| $T$ | 200 | dt | Total timesteps |
| $dt$ | 50 | ms | Timestep |
| $\tau_b$ | 1.31 | s | “Backwards” kernel time constant |
| $\tau_f$ | 0.69 | s | “Forwards” kernel time constant |
| $\lambda$ | 1 | - | Weight constant |
| $\Delta t$ | 5 | s | Time window of integration around plateau |
| $\gamma$ | 10 | - | Constant input, to go from $\Delta W$ to $\Delta V$ (Figure 1) |
| $\sigma$ | 0.075 | s | Standard deviation of input Gaussians |
| $\theta$ | 8 | - | Threshold for plateau event (Figure 2) |
| $N_{input}$ | 200 | - | Number of input neurons |
| $N_{output}$ | 81 | - | Number of output neurons |
| $\eta$ | 0.95 | - | Learning rate (Figure 2) |
| $\sigma_n$ | $\mathcal{N}(0, 1/35)$ | - | Noise, output neuron activity |
| $L$ | 100 | cm | Box side length |
| $\sigma_w$ | $\mathcal{N}(0, 1)$ | cm | Random walk updates (2D) |
| $\vartheta$ | 0.15 | - | Threshold for plotting neural activity (Figure 2i) |
| $N_{seed}$ | 81 | - | Number of seed neurons |
| $N_{visible}$ | 100 | - | Number of visible neurons |
| $\theta$ | .6 | - | Threshold for plateau event (Figure 3) |
| $\eta$ | 0.3 | - | Learning rate (Figure 3) |
| $W^{in}$ | $\mathbb{I}$ | - | Input weights (Figure 3) |
| $C$ | 2.71 | - | Constant for gating term |
| $N_{input}$ | 100 | - | Number of input neurons (Figure 4) |
| $N_{inter}$ | 10 | - | Number of hidden neurons (Figure 4, fixation task only) |
| $N_{output}$ | 1, 2 | - | Number of output units (Figure 4, place cell task, fixation task) |
| $T$ | 100 | dt | Total timesteps (Figure 5) |
| $\tau_{net}$ | 10 | dt | Network time constant (Figure 5) |
| $N_{input}$ | 2 | - | Number of input neurons (Figure 5) |
| $N_{rec}$ | 100 | - | Number of recurrent neurons (Figure 5) |
| $N_{output}$ | 2 | - | Number of output neurons (Figure 5) |
| $g$ | .85 | - | Gain, recurrent weight initialization (Figure 5) |
| $\eta$ | 0.01 | - | Learning rate, ADAM (Figure 5) |
| $\beta_1$ | 0.9 | - | Decay rate for first moment estimate in output weights, ADAM (Figure 5) |
| $\beta_2$ | 0.99 | - | Decay rate for second moment estimate in output weights, ADAM (Figure 5) |
| $\beta_{1,rec}$ | 0.999 | - | Decay rate for first moment estimate in recurrent weights, ADAM (Figure 5) |
| $\beta_{2,rec}$ | 0.9999 | - | Decay rate for second moment estimate in recurrent weights, ADAM (Figure 5) |
| $\epsilon$ | $10^{-8}$ | - | Constant to avoid division by zero, ADAM (Figure 5) |

### Generalized Learning Rule for Behavioral Timescale Plasticity

To begin our derivation, we consider the simple case of a single postsynaptic neuron which triggers an instantaneous plateau event, and a single presynaptic neuron which fires a spike (**Figure 1a**). We assume the postsynaptic plateau updates weight *W* via some function, *W*_*kernel*_, which depends on the timing of the presynaptic spike relative to the plateau:

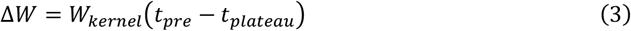

Specifically, we choose a *W*_*kernel*_ such that the application of our learning rule matches observed plasticity following application of a single plateau and bursting inputs in vitro (**Figure 1d**)^2^. The specific form of the weight kernel is:

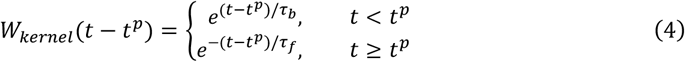

Where *t*^*p*^ is the time of the plateau, and *τ*_*b*_ and *τ*_*b*_ are “backward” and “forward” time constants.

Next, we relax our previous assumption that there is a single presynaptic neuron which fires a single spike, instead describing the continuous activity of presynaptic neuron *j* at time *t* as *x*_*j*_(*t*) (**Figure 1b**). Now, the change in weights following a single plateau depends on the integrated presynaptic activity across a temporal window Δ*t* relative to the time of the plateau):

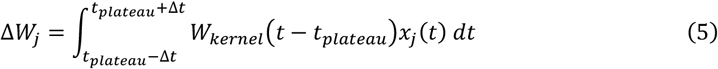

Note that if *x*_*j*_(*t*) is taken to be a delta function *δ*(*t* − *t*_*pre*_), Equation 5 reduces to Equation 3. Further, the asymmetric offset of observed plasticity in **Figure 1e** is a direct consequence of the shape of *W*_*kernel*_ (Equation 4). For some intuition on why this is the case, notice that Equation 5 is equivalent to a cross-correlation, so we can imagine “sliding” or “smearing” *W*_*kernel*_ across the input *x*_*j*_(*t*) to get a given horizontal slice of **Figure 1e**.

To match experimental data showing that the amplitude of the formed field depends on the initial membrane voltage of the postsynaptic cell^1^, we add in a dependence on the synaptic strength prior to the plateau event, leading to the equation:

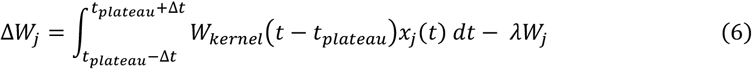

where the weight dependence is parametrized by λ. Note since Δ*W* is only applied when a plateau occurs, −*λW* is not a continuous weight decay.

Finally, we want to consider the case for which there are multiple postsynaptic neurons, each of which may have multiple plateaus. So, we introduce *P*_*i*_(*t*), a function representing the post-synaptic plateau potential at time *t* for neuron *i*. Now, the change in weights depends on an integral over the presynaptic activity, as well as an integral over any post-synaptic plateaus (**Figure 1c**):

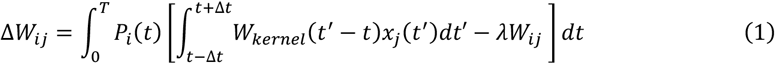

in which weight *W*_*ij*_ is updated after each trial according to the presence of *P*_*i*_(*t*). Note that if *P*_*i*_(*t*) is taken to be a delta function *δ*(*t* − *t*_*plateau*_), Equation 1 reduces to Equation 6. Since Equation 1 is derived from various degrees of generalization (Equations 3, 5, and 6), we call this equation “generalized BTSP”.

### Unsupervised Feed-Forward Task

Notice in Equation 6, that if we take the temporal kernel *W*_*kernel*_(*t* − *t*_*plateau*_) to be the delta function *δ*(*t* − *t*_*plateau*_), the integral over time goes away and we get the following expression:

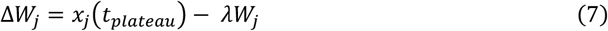

So, in this approximation, upon each plateau event, the weights would move towards a fixed Point 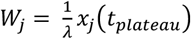, akin to classical conceptions of “competitive learning”^17–20^.

For the 1-D unsupervised feed-forward task in **Figure 2b-e**, we assume an animal is running along a 1-D treadmill at constant velocity *β*, i.e. the animal’s position *u*(*t*) = *βt*. The external sensory input is modeled in the form of stereotypical 1-D tuning curves with added noise:

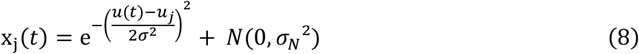

where j indexes over *N* total inputs, and *u*_*j*_ are the locations (or equivalently, times) of the tuning curve centers, which have standard deviation σ. Zero-mean Gaussian noise is added, with standard deviation *σ*_*N*_. These inputs are connected to an output *y*_*i*_(*t*) by feed-forward weights *W*_*ij*_:

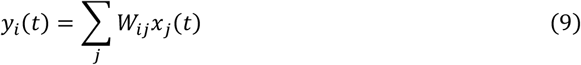

which are learned via gBTSP. We used the following rule to trigger a plateau:

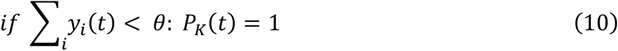

Where *θ* is a firing rate threshold, and *K* is a random index from 1 to N. A single plateau can drive the network above the threshold *θ* (at a given time). Weights were initialized at zero and the network was trained on 100 laps.

For the 2-D unsupervised feed-forward task in **Figure 2f-h**, we assume an animal begins at a random location inside a 2-D box, (*u*_0_, *v*_0_) and takes *T* steps of a random walk along a trajectory (*u*(*t*), *v*(*t*)). The external sensory input is modeled in the form of stereotypical 2-D tuning curves with added noise:

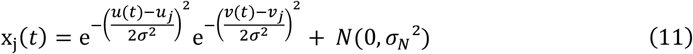

where j indexes over *N*^2^ total inputs, and (*u*_*j*_, *v*_*j*_) are the 2-D locations of the tuning curve centers, which have standard deviation σ. Zero-mean Gaussian noise is added, with standard deviation *σ*_*N*_. These inputs are connected to an output *y*_*i*_(*t*) by feed-forward weights *W*_*ij*_ (Equation 9), which are learned via gBTSP, just as in the 1-D case.

### Unsupervised Recurrent Task

For the unsupervised recurrent task in **Figure 3**, we assume an animal is running along a 1-D treadmill at constant velocity *β*, i.e. the animal’s position *u*(*t*) = *βt*. The external sensory input is modeled the same as for the unsupervised feed-forward task (Equation 8). There are two populations of neurons, “visible” neurons *x*_*j*_(*t*) and “seed” neurons *s*_*i*_(*t*). These populations connect recurrently to each other, but not amongst themselves. Visible neurons receive external input *o*_*k*_(*t*), and only the seed neurons eligible for plateaus (**Figure 3a**). The activity of these two populations is governed by the following equations:

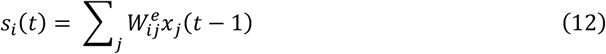

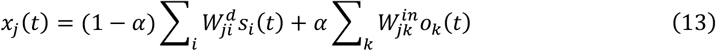

where 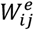 are “encoding” weights, 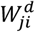 are “decoding” weights, and 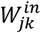 are input weights. *α* is a gating variable governed by the norm of the external input: when external input is high, recurrent input is low, and vice versa:

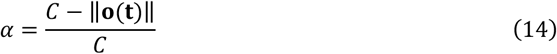

Where C is a constant and ‖**o(t)**‖ is the norm of the external input. In the limit of no external input (*α* = 0), our equation for the visible neurons reduces to:

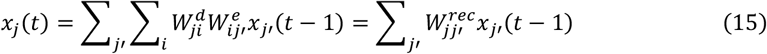

where 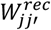 is the low-rank recurrence, 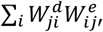 of the visible neurons. However, in the limit of large external input, our network becomes effectively feed-forward:

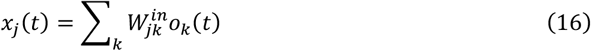

This dual nature of the network allows us to take advantage of recurrent computation in the low-input phase, while making use of unsupervised, feed-forward gBTSP in the high-input phase. Encoding weights are learned via the same competitive learning algorithm as the feed-forward case (Equation 10). The decoder weights are set to be the transpose of the encoder weights. Following a single lap of training, a test phase was conducted, whereby inputs were only shown for one timestep before being removed. The inputs were shown at times 0, 2, 4, 6, and 8 seconds.

### Supervised Feed-Forward Task

In order to apply plateaus in the supervised context, we derive an expression for *P*(*t*) which minimizes a given error/loss. Assuming some target output *ŷ*(*t*), and a loss function ℒ (here we choose a mean squared error loss).

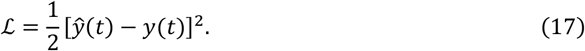

We can compare our gBTSP weight update

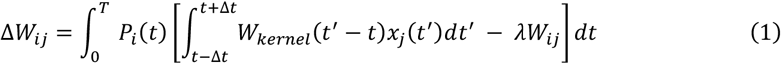

to that of simple backpropagation (for a single layer, this is just the delta rule):

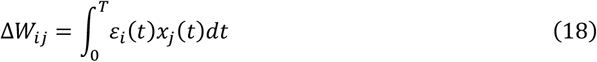

Where the error term *ε*_*i*_(*t*) = *ŷ*_*i*_(*t*) − *y*_*i*_(*t*). By setting these two equations to be equal, we can find an expression for the function *P*_*i*_(*t*):

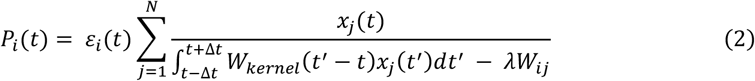

For the first task, we consider a shallow feed-forward network (Equation 9). We choose a target function *ŷ* (*t*) that is a putative place field, modeled as a Gaussian bump centered at a specific location in the environment:

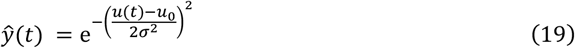

Where *u*(*t*) is the animal’s position, and *u*_0_ is the location of the tuning curve center, which has standard deviation σ.

For the navigation task, the network has three layers (input, hidden, output), for which only the hidden neurons can receive plateaus, i.e. only the input to hidden weights *W*_*jk*_ are trainable:

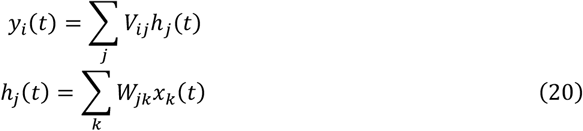

The 2D target trajectory for the task is:

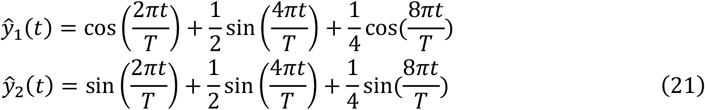

We train the network (Equations 1 and 2) for 100 trials on both tasks.

### Supervised Recurrent Task

For our recurrent task, a network of hidden units with activation *h*_*j*_(*t*) is recurrently connected via weights 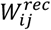. The dynamics of the hidden units are governed by the following:

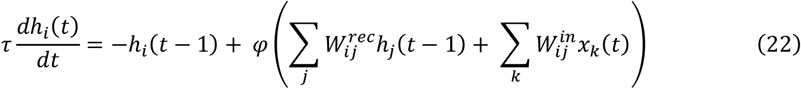

here φ is a non-linear function (i.e. tanh) of the recurrent inputs, and these activations are initialized at *h*_0_. Input *x*_*k*_(*t*) is projected to the network via input weights 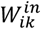. We initialize the internal weights as a random Gaussian matrix with a gain factor *g*. These hidden units project to output *y*(*t*) via weights 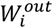:

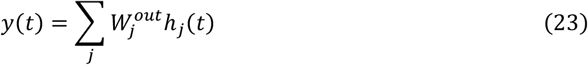

To find *P*_*i*_(*t*) which minimizes the error in a recurrent network, we again compare our gBTSP rule

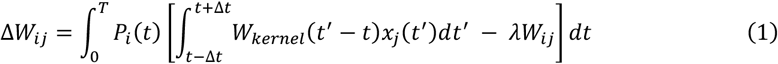

to the full backpropagation through time update: (for a full derivation, see Murray, 2019^13^):

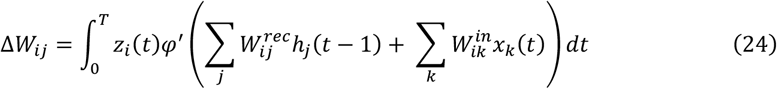

where φ′ is the derivative of our activation function, and the Lagrange multiplier *z*_*i*_(*t*) is equal to:

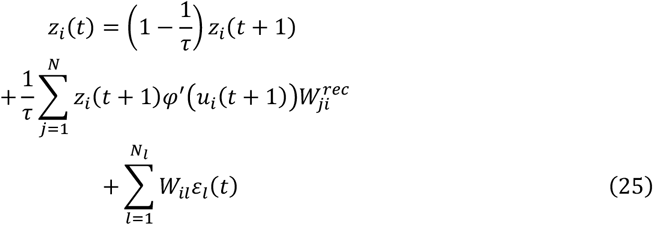

where *u*_*i*_(*t*) is our total input current to the unit, i.e.,

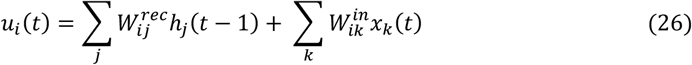

The Lagrange multiplier *z*_*i*_(*t*) is calculated in the “backwards” phase, by starting with the terminal value, *z*_*i*_(*T*), and working back to *z*_*i*_(0). *z*_*i*_(*T*) takes the form:

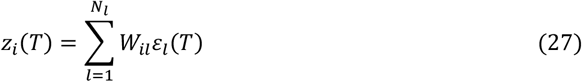

By setting Equation 1 and Equation 24 to be equal, we solve for the plateau function and get the following expression:

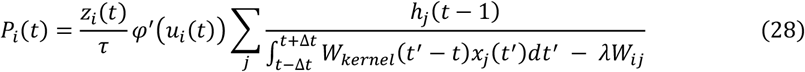

The delayed non-match to sample task consists of two inputs, representing odor A and odor B, and two outputs: one representing licking probability and the other which represents a ramping temporal component. This component encourages the network to develop sequential internal representations^47^. For each trial, a random odor combination (AA,AB,BA,BB) was selected. The first odor input was presented for the first 1s of the trial, and the second odor input was present between 7-8s. No odor inputs were given in the delay period. For non-matching pairs, the target for licking probability was 1 for all timesteps after 8 seconds, and 0 otherwise. For matching pairs, the target licking probability was always 0. Output weights were trained using the delta rule, while recurrent weights were trained using our gBTSP update (Equation 1), after selecting plateaus (Equation 28). The network was trained for 50,000 trials, with the gBTSP update (Equation 1) passed through a momentum-based optimizer (ADAM^46^) to avoid critical instabilities (see Table 1 for parameters).

### Constraints on Few-Shot Learning

In the case of the shallow network, inputs *x*_*j*_(*t*) project to output *y*_*i*_(*t*) = *∑*_*j*_ *W*_*ij*_*x*_*j*_(*t*). (**Figure 6b**). “Plateaus” of size Δ*x* occur at the inputs, and the loss is the mean squared error between output **y** and target *ŷ*. Solving for the Hessian (local curvature), we find:

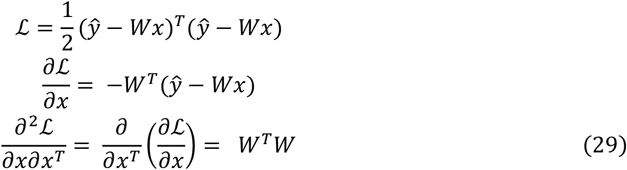

If we consider a deep feed-forward network with layers *l* and layer specific weights *W*_*l*_ (**Figure 6c**), the expression for the Hessian becomes more complicated:

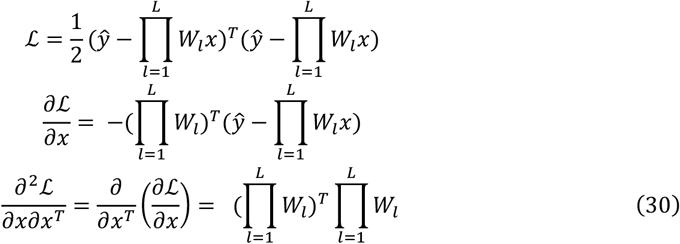

For recurrent networks (**Figure 6d**), the Hessian in a recurrent network depends on the *T*th product of *W*, which also leads to exploding and vanishing contributions from an update Δ*x*:

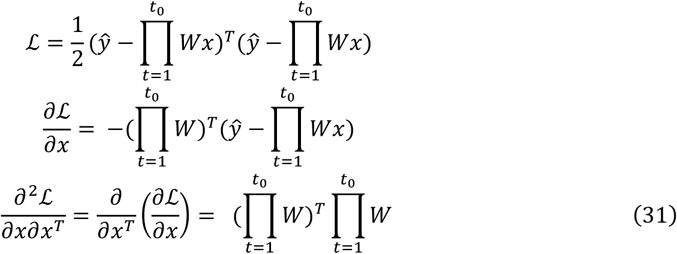

These conditions set fundamental limits on both the architectures and the tasks for which deep or recurrent artificial networks can support rapid changes in single-unit activity.

## Acknowledgements

This work was supported by BBSRC (BB/N013956/1), Wellcome Trust (200790/Z/16/Z), the Simons Foundation (564408), EPSRC (EP/R035806/1 and EP/X029336/1) and ERC-UKRI (EP/Y027841/1).

## Author Contributions

I.C., C.C. and R.P.C. conceived and designed the model. I.C. developed and performed the simulations. I.C., C.C. and R.P.C. wrote the manuscript.

## Supplemental Figures

**Supplemental Figure 1.**
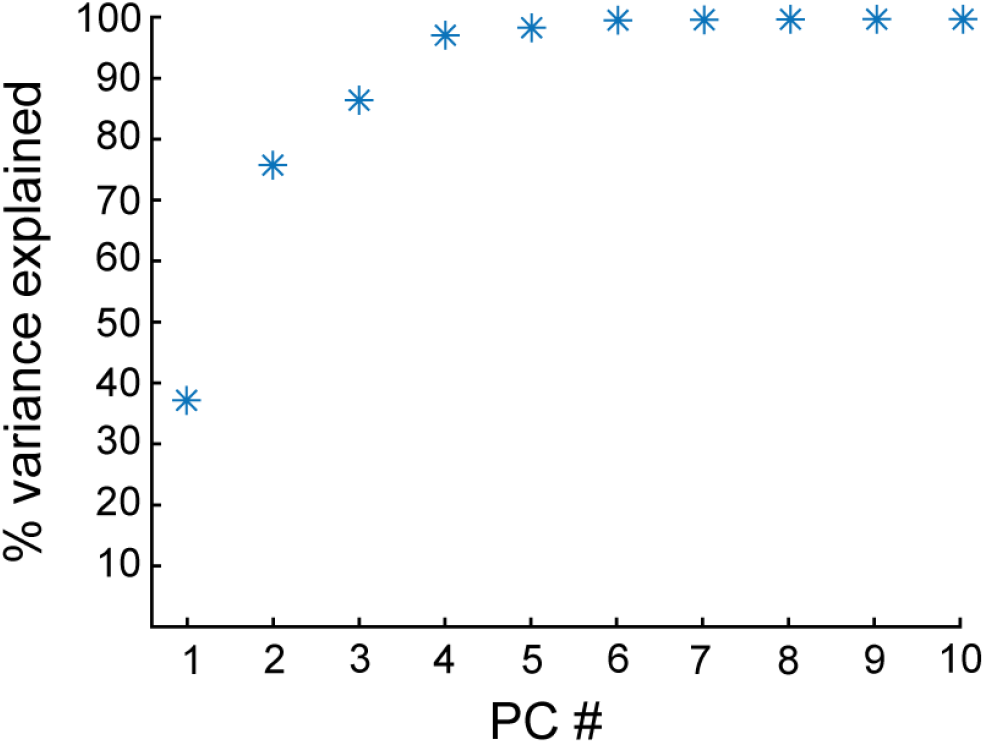
Variance explained by principal components in recurrent attractor network. Cumulative sum of variance explained by the first ten principal components of the effective recurrent weight matrix, **W**^**rec**^ = **W**^**e**^**W**^**d**^, after competitive learning via gBTSP. The first two components are shown in Figure 3c.

**Supplemental Figure 2.**
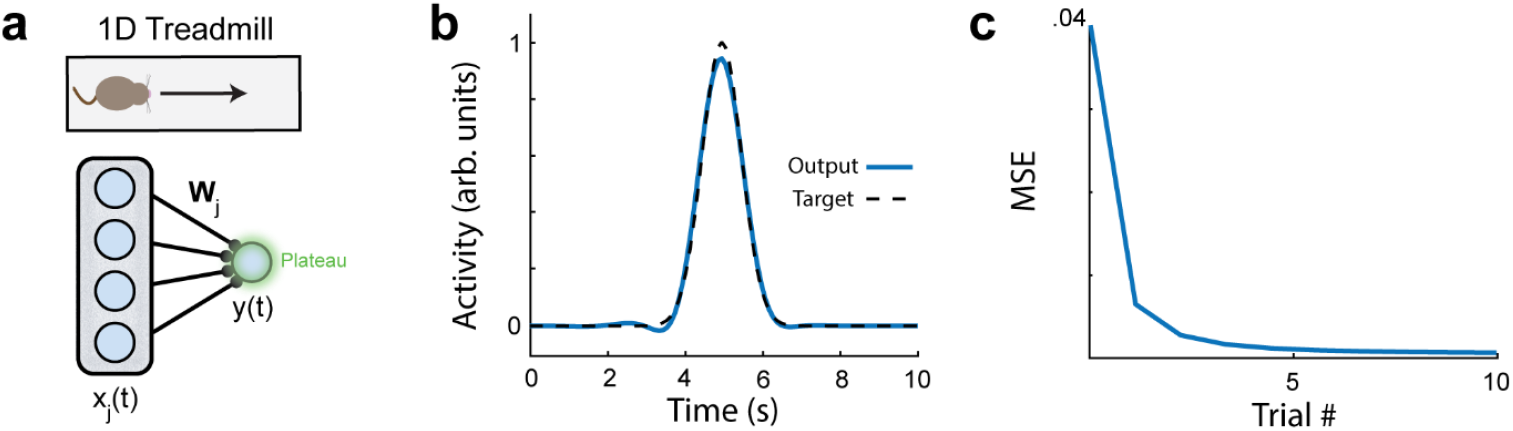
Mean squared error for supervised feed-forward tasks. **a)** Feed-forward network with inputs *x*_*j*_(*t*) project to output *y*(*t*) via weights *W*_*j*_. The inputs *x*_*j*_(*t*) are spatially selective and represent an animal running along a 1D track. **b)** A unimodal gaussian target function (dotted black line) and the trained output (blue line) after the first 10 trials of gBTSP training. **c)** The mean squared error over the first 10 trials of training.

**Supplemental Figure 3.**
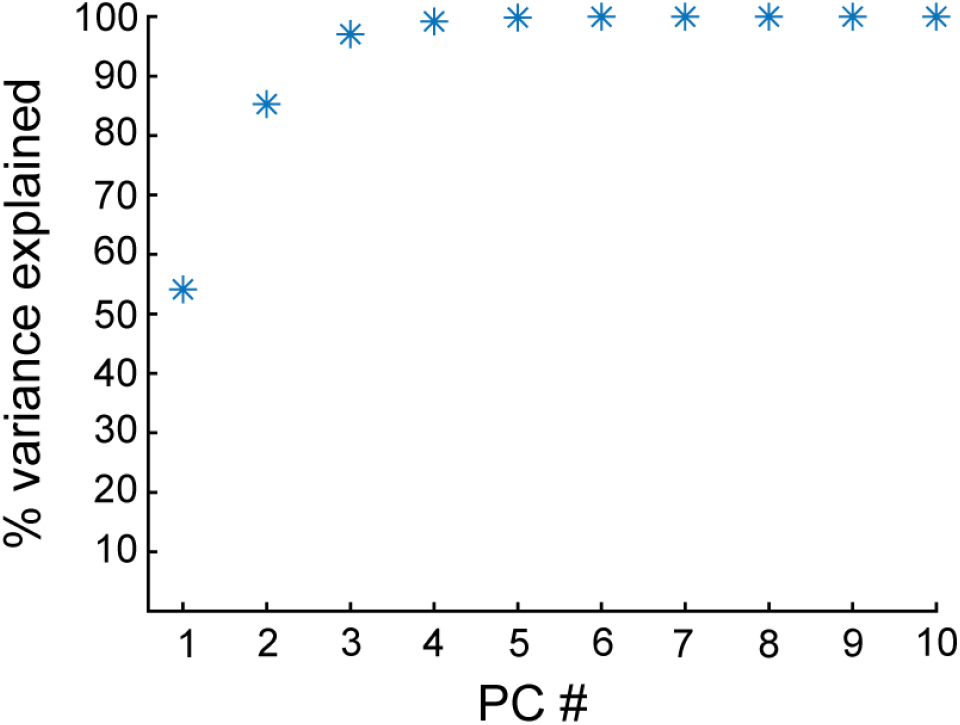
Variance explained by principal components in DNMS task. Cumulative sum of variance explained by the first ten principal components of network activity in the DNMS task. The first three components are shown in Figure 5e.

**Supplemental Figure 4.**
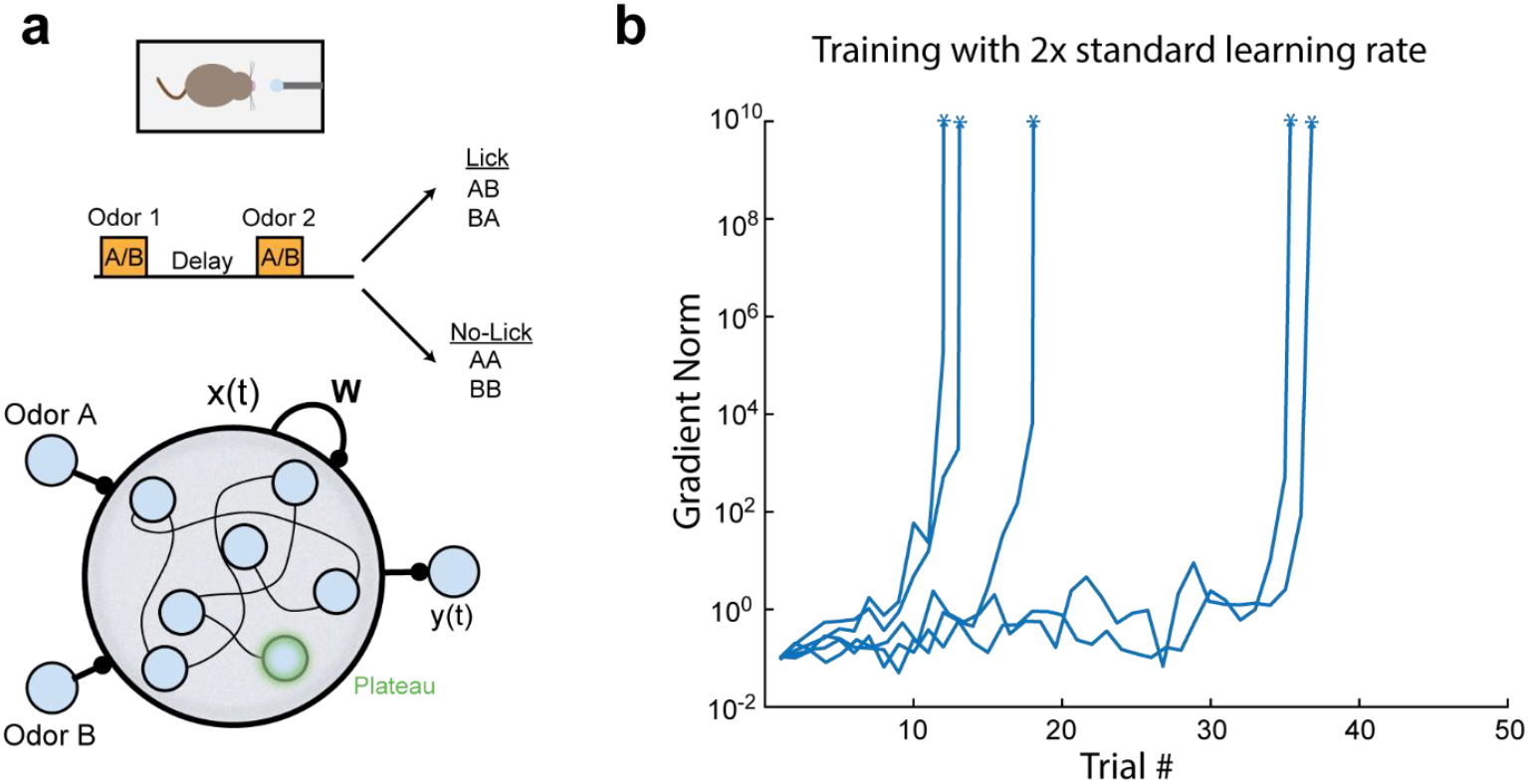
Large learning rates lead to instability in gBTSP in recurrent networks. a) Simulated agents are trained on a delayed-non-match-to-sample (DNMS) task where they m ust distinguish between sequential “odor” pairs. The agent must learn to lick for non-matching sequences (AB, BA), and refrain from licking for matching sequences (AA,BB). Bottom, the model consists of a recurrent network with two odor inputs, hidden activities **x(t)** and recurrent weights **W**. The hidden units project to an output **y(t)** via weights **V**. Recurrent weights **W** are trained via plateaus occurring in the hidden units according to our gBTSP algorithm (see Methods). Output weights are trained via the delta rule. **b)** Training using a learning rate double that used in Figure 6 (see also Table 1) results in gradient explosion. First five random seeded runs shown. Y-axis is the norm of the gradient of the recurrent weights and is truncated at a value of 10^10^ for visual clarity.

